# The fMRI global signal and its association with the signal from cranial bone

**DOI:** 10.1101/2024.03.27.587003

**Authors:** Daniel Huber, Luna Rabl, Chiara Orsini, Karin Labek, Roberto Viviani

## Abstract

The nature of the global signal, i.e. the average signal from sequential functional imaging scans of the brain or the cortex, is not well understood, but is thought to include vascular and neural components. Using resting state data, we report on the strong association between the global signal and the average signal from the part of the volume that includes the cranial bone and subdural vessels and venous collectors, separated from each other and the subdural space by multispectral segmentation procedures. While subdural vessels carried a signal with a phase delay relative to the cortex, the association with the cortical signal was strongest in the parts of the scan corresponding to the laminae of the cranial bone, reaching 80% shared variance in some individuals. These findings suggest that in resting state data vascular components may play a prominent role in the genesis of fluctuations of the global signal. Evidence from other studies on the existence of neural sources of the global signal suggests that it may reflect the action of multiple mechanisms (including cerebrovascular reactivity and autonomic control) concurrently acting to regulate global cerebral perfusion.

## Introduction

The existence of fMRI signals not originating from neural activity, such as those arising from respiration, heartbeat, and their modulation (Wise et al. 2004; Birn et al. 2006, 2008; Shmueli et al. 2007; Chang et al. 2009) has been long recognized (“physiological noise”). In rest connectivity studies, but also in task-based fMRI (Aguirre et al. 1998), signals of non-neural origin and effects of head movements have been considered a potential confounder to be removed (Power et al. 2012; Murphy et al. 2013). Several approaches have been proposed in the literature to this effect, including the use of the mean signal from white matter or the ventricles as confounding covariates (Behzadi et al. 2007; Power et al. 2014).

Increasing attention is recently being paid to the “global signal”, i.e. the average signal arising from the brain (Zarahn et al. 1997; Liu et al. 2017). It has been shown (Power et al. 2017) that this signal is practically identical to the mean signal from the cortex (here, we consider the mean signal from cerebral cortex and the expression “global signal” to be synonymous, the former being a good estimate of the latter). Like the mean signal from other tissue classes such as white matter or the ventricles, the global signal has been used as a confounder in connectivity fMRI analyses to control for signal of non-neural origin (for review, see Liu et al. 2017). Furthermore, the global signal has also been shown to be associated with individual differences of clinical nature, leading to criticism of its use as a confounder (Gotts et al. 2013; Hahamy et al. 2014). To date, the nature of the global signal is not fully understood (Liu et al. 2017) but it appears to include the effect of vascular reactivity to changes in the partial volume of CO_2_, such as those following brief respiratory events (Power et al. 2017). Other studies, however, have provided evidence of central neurogenic control of the global signal (Liu et al. 2018; Turchi et al. 2018).

A major issue concerning the global signal is the consideration that, emerging prevalently from grey matter, it may include signal of neural origin. In the present work, we describe the association of the mean signal from cerebral cortex with the average signal from the “cranial bone segment” in fMRI data of a public dataset of images acquired at rest (Mendes et al. 2019), where the abundant accompanying structural imaging allowed us to refine the segmentation of the bone tissue class. Our rationale for looking at the mean signal from bone was that, while of course vascularized, bone is not expected to include neural tissue. Hence, we thought of it as a possible selective assay of generic peripheral vascular confounders and/or effects of head movements.

The quotation marks around “cranial bone” emphasize that this is a concept borne out of segmentation procedures in MRI practice and not an anatomical fact (the same qualification applies to other anatomical terms, such as “grey” or “white matter”). Nevertheless, the correspondence between segmentation output and tissue properties can be enhanced by including multiple sources of signal (multispectral segmentation, Lambert et al. 2013, Viviani et al. 2017). Here, we used this technique to increase the specificity of the bone segment and identified additional tissue classes that corresponded spatially to dural venous sinuses and the bone itself and that appeared to provide dissociable contributions to the mean signal. These sub-segments are characterized by low signal intensities in EPI images but are not completely devoid of signal.

The association between the mean cortical and bone signals (but not between cortex and soft tissue) is apparent without adopting specific techniques but is clearest in the frequency range typical of connectivity studies (0.008-0.1 Hz), which allows to exclude shared signal confounders such as the low-frequency scanner drift. In the analyses of the Results section, we varied the method settings to verify the robustness of the finding. We found that the association did not depend on whether the signal was extracted from volumes in registered or in native space, whether head motion was conspicuous or not, or whether parts of the head or neck were included or excluded from the bone segment.

To gain insights into its possible mechanisms, a second set of analyses was conducted with multispectral segmentation techniques (Lambert et al. 2013, Viviani et al. 2017) to better separate the bone from intracranial space and subclassify the bone tissue class into separate sub-components. An important aim was to identify venous collectors (dural sinuses or meningeal veins), which are known to contain signal that follows that of cortex with a phase delay (Tong and Frederick 2010; Liu et al. 2017; Mitra et al. 2014) and run in the subdural space, and separate them from bone, and characterize the contribution of head motion. In the Discussion, we review possible artefactual and physiological explanations of our findings, emphasizing the importance of understanding the role of vascular signal in functional imaging data to identify sources of individual differences.

## Results

### Standard segmentation

In this first analysis, we considered data and segmentation conducted with the default settings of the SPM package (Ashburner and Friston 2005), which would be representative of the data used in a standard functional imaging analysis (data realigned and registered to MNI template). However, the data from which the mean segment signal was extracted were not smoothed to avoid contamination from contiguous tissues. We also eroded the tissue classes to minimize partial volume effects of voxel adjacent to neighbouring tissue classes. Erosion was 2 voxels except for grey matter, which was eroded by 1 voxel (Figure 1A shows the resulting classification of voxels into tissue classes and their spatial separation). In all correlations, head motion estimated from realignment, its first derivatives and the quadratic terms (24 parameters) were partialled out.

**Figure 1.**
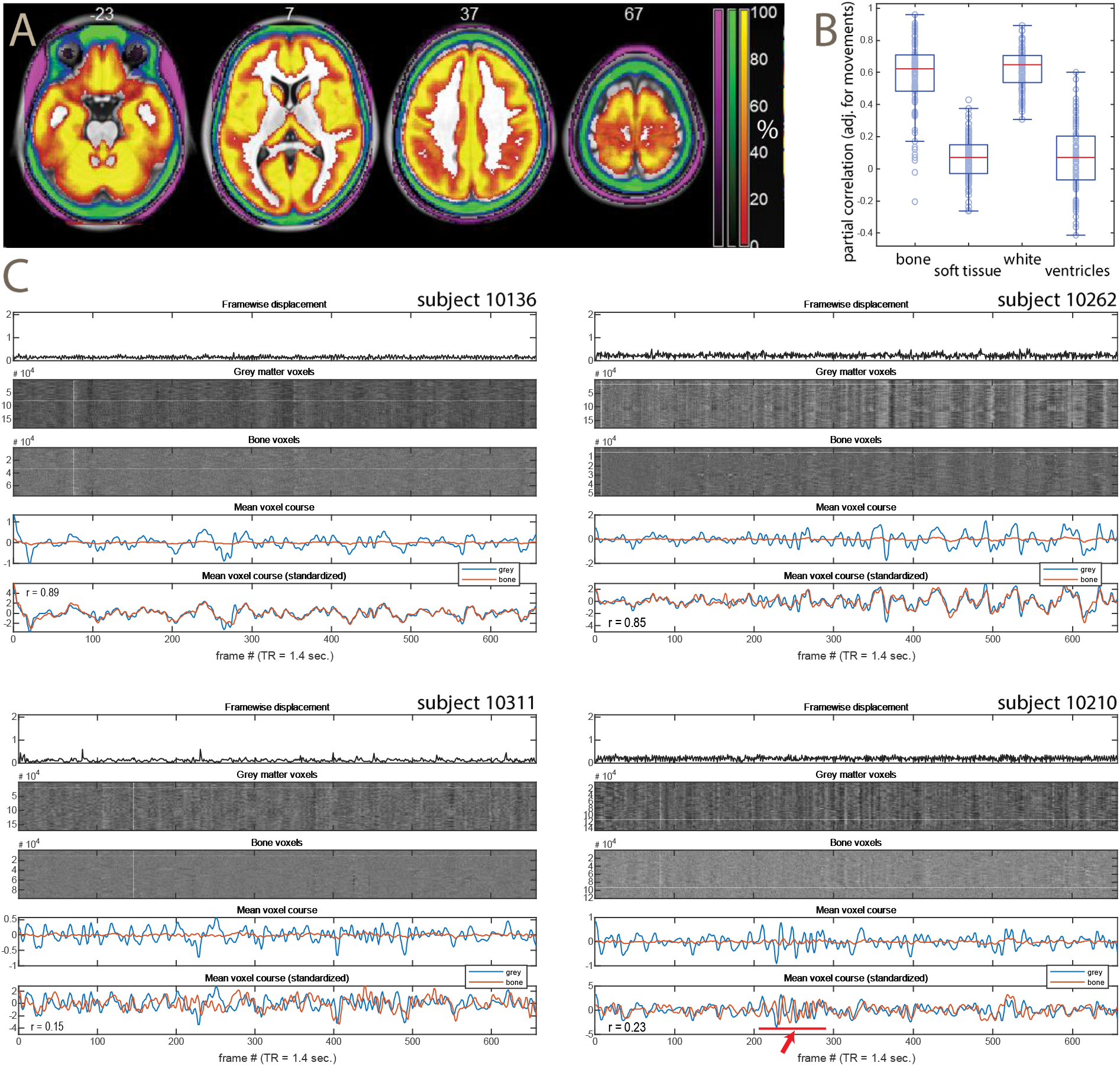
A: summary maps of the grey, bone, and soft tissue segmentations (percent of voxels classified in the segment), showing the separation between these tissue classes in the analysis. Pure yellow indicates that the voxel was classified in all participants as grey, full green as bone, violet as soft tissue. Because of the individual variation in the classification of voxels at tissue class boundaries, the separation is larger in the individual segmentations (i.e. the figure shows the minimal bound of the separation). B: boxplot of partial correlations between average grey matter and bone or soft tissue signal courses (partialled for movement covariates, their first derivative, and their quadratic terms). C: example of signal courses from four subjects (at the top, second and third best correlations; at the bottom, second and third worse correlations between grey matter and bone segments). In each subject, from top to bottom: the framewise displacement, the intensity of the signal in the grey matter segment, in the bone segment (both smoothed FWHM 8mm), the average signal in these two segments (without spatial smoothing, band-pass filtered, de-meaned), the average signal in these two segments (without spatial smoothing, band-pass filtered, standardized).

Figure 1B shows boxplots of the correlations between the mean grey matter signal course and that of bone (left), soft tissue, white matter, and ventricles (right). One can see that the correlation between bone and grey matter (first box on the left) averages about 0.6, although a few subjects do not show correlation, and others show a correlation as high as 0.9. This implies that in these individuals about 80% of the variance in the global signal from grey matter and bone was shared. In contrast, the correlation between grey matter and soft tissue (second from left) is not substantial. The correlation of the bone segment is similar to that of white matter (third from left), while the ventricles show a smaller correlation.

Figure 1C shows the signal course in four individuals, chosen to be representative of individuals with high (top row) and low correlation (bottom row; we chose the second and third best or worse, to avoid the possible outliers at the extremes). In each displayed individual, the sub-panel at the top shows the framewise displacement. One can see that none of the individuals displayed in this Figure showed large movements. Nevertheless, there was a very large correlation between the grey and bone mean signal courses in the two individuals in the top row.

Below framewise displacement, two sub-panels show “heatmaps” (Power 2017) of the signal in the grey and bone matter, i.e. a plot of the signal intensity of all voxels within the segment. Following Power (2017), the heatmap data (but only those) were smoothed prior to being displayed, as otherwise the tendency of the mean signal does not emerge visually. One can see that there were considerable modulations of the signal even in the absence of large movements, more marked in subject 10262 on the top right. One can also see these broad modulations in the bone segment voxels, even if the contrast is much lower. As discussed in Power et al. (2017), these modulations may follow larger head movements indexing respiratory events, but they also commonly appear in the heatmaps of grey matter even in the absence of movements, as is the case in these participants.

The two bottom panels show the de-meaned average signal course of the grey (blue) and bone (red) segments in each subject. The segments are the same as those of the heatmaps, but the signal was not spatially smoothed prior to computing the course and was band-pass filtered at 0.1 and 0.0078Hz (for this reason, some high-frequency oscillations in the heatmaps may not be visible in the average signal course). In the first of these two panes, one can see the signal of bone having much lower amplitude than the other. This also reflects the lower signal intensity of bone in EPI images, and accounts for the lower contrast in the heatmaps of bone relative to grey matter voxels. In the second sub-panel at the bottom, the signal was standardized to unit variance in both grey and bone signal courses (an operation carried out internally by the computation of the correlation). Here, one can see how the average signal from bone closely follows that of grey matter.

In the two subjects at the bottom the correlation between bone and grey matter was absent. In subject 10311 on the left, there was less average signal to begin with, relative to the mean image intensity (the range of the y axis in the de-meaned average signal course is smaller than in the other subjects). In subject 10210 on the right, one can see that the signal in bone was delayed relative to the grey matter signal. This is particularly apparent in the portions where the amplitude of the signal was larger, such as the part highlighted by the red arrow and bar. This suggests that the signal from grey matter was present also in this subject, but the delay led to a decrement of formal correlation values.

A question naturally arising from these findings is the extent of the association between bone and grey matter when the signal is extracted from the images in native space. These images are the closest to those delivered by the scanner (the only applied preprocessing step was realignment). The boxplots showing partial correlations between grey matter and bone and, for comparison, soft tissue, are shown in Figure 2.

**Figure 2.**
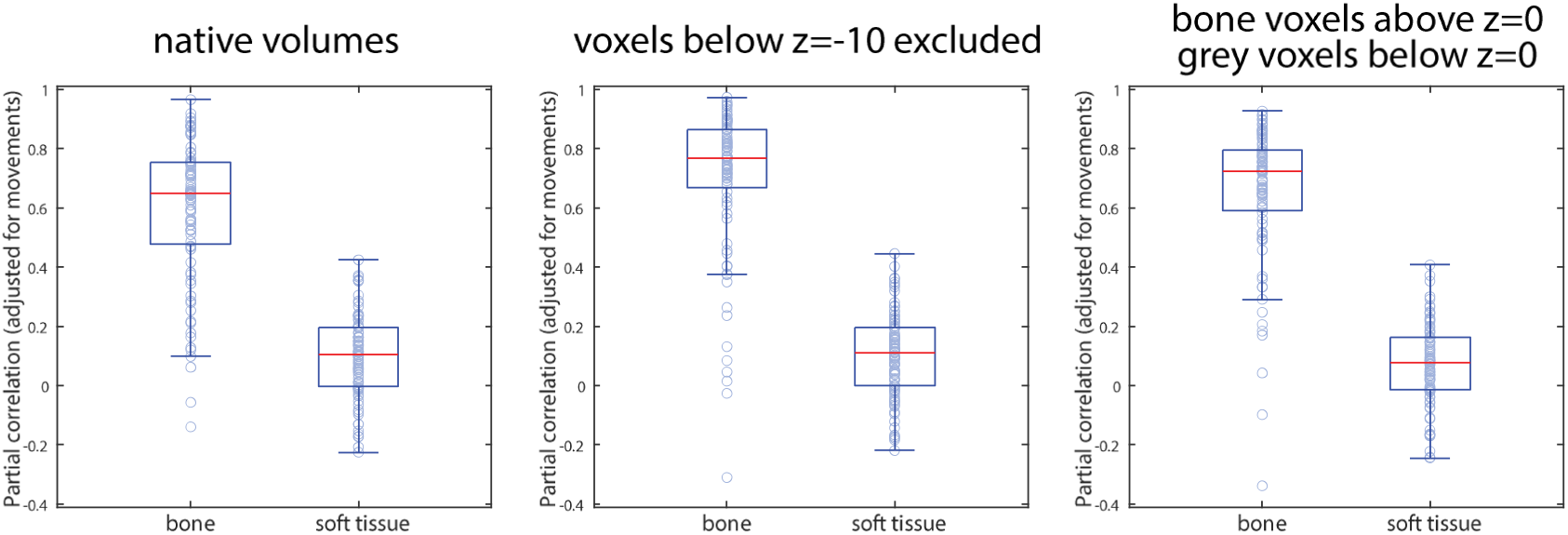
Boxplots of partial correlations between average grey matter and bone or soft tissue signal courses (adjusted for movement covariates, their first derivative, and their quadratic terms) from volumes in native space, with masks to exclude contributions from head portions.

One possible explanation of the association between bone and grey matter is given by aliased heartbeat signal in vessels at the base of the brain and head, which may be present in both bone and grey matter. To investigate this possible source of the correlation, we repeated the analysis on the volumes in native space while excluding all voxels below *z* = −10 (after applying the reverse registration transformations to the masks from MNI space into native space), to avoid recruiting large vessels at the base of the brain (Viviani 2016). The results are shown in the boxplot in the middle of Figure 2. One can see that excluding these voxels led to an increase in the correlations, against the assumption of a confounded aliased heartbeat signal. One can see here that the majority of individuals had a correlation of 0.8 or larger, implying that 60-80% of shared variance between the mean signal from cortex and bone was common in the sample.

Another possible explanation of the correlation findings is the effect of Gibbs artefacts contaminating signal in the bone segment from signal from cortex. If this is the case, one would expect the correlation to depend on the signal being collected in the same slice (which would be here transversal slices due to the acquisition modality). The right boxplot in Figure 2 shows the average signal from voxels in the bone segment above *z* = 0 in MNI space, while the average signal from grey matter was extracted from voxels below this plane. Again, one can see that the correlation, while less marked than when acquired from the whole volumes, is still very much larger than for soft tissue.

### Multispectral segmentation

In the second analysis, the segmentation was conducted using a multispectral approach with five input channels (mean functional image and four structural images). Because MRI images acquired with different techniques provide differing contrast between tissues, combining the information from many image types may allow separating tissue classes that have the same intensity in the functional images of the previous analyses. Our purpose was to improve the specificity of the bone segment, especially for what concerns venous collectors which, as mentioned in the Introduction, may convey signal from cortex. To this end, we modelled vessels as its own separate tissue class. The signal intensity of the bone segment was modelled by a mixture of two Gaussians. Because the resulting tissue segments are much thinner than those of the default segmentation, we ensured the specificity of the voxels included in the segment by thresholding the probability maps computed by the segmentation at 0.75, instead of using erosion (which would have led to either few remaining voxels and/or highly varying voxel numbers within the sample). This was especially necessary for vessels and the bone sub-segments since their dimensions spanned only few voxels (or even one) in most regions. Grey matter and soft tissue were eroded by 1 voxel. Unless otherwise mentioned, all images and results presented in this section refer to native space data. Figure 3 shows sections of the segmentation from two representative individuals, paired with the structural T2-weighted images.

**Figure 3.**
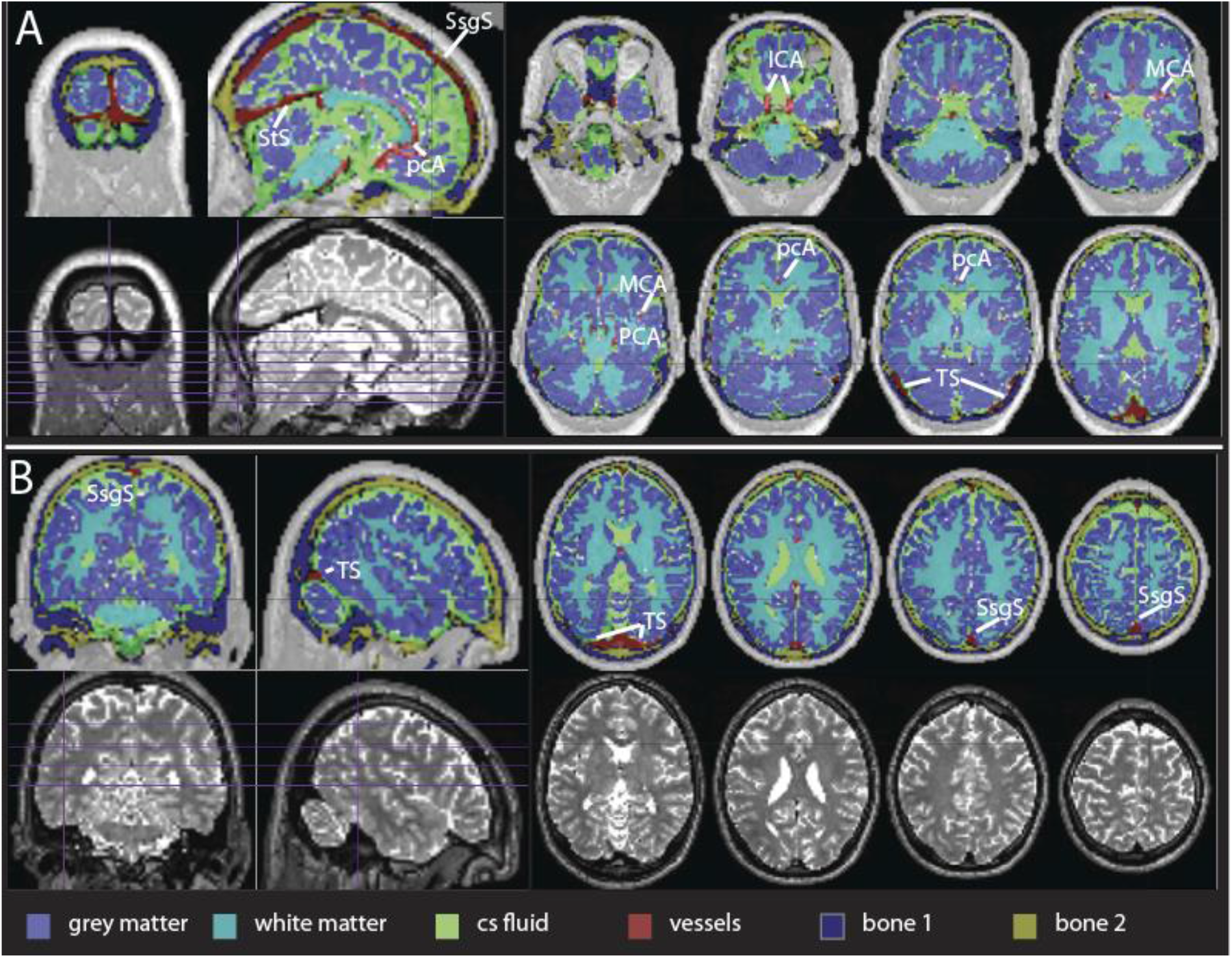
Co-registered T2-weighted images with overlay of assigned tissue classes. Part A presents slices chosen to illustrate the segmentation of the vessels (comprising a mixture of arterial and venous), while part B focuses on the separation of the bone sub-segments. Comparison with the T2-weighted images (bottom row) shows this sub-segmentation to rely on brighter signal from the central part of the cranial bone. SsgS: superior sagittal sinus; StS: straight sinus; TS: transversal sinus; pcA: pericallosal artery; ICA: internal carotid artery; MCA: middle cerebral artery; PCA: posterior cerebral artery.

One feature of this segmentation that is worth focusing on is the capacity to identify venous collectors as a separate tissue class. In Figure 3A, the sections demonstrate the separation of the vessels (red) from the surrounding tissues. The large dural venous sinuses are clearly visible along with some smaller vessels. The success of vessel segmentation was visually checked by directly overlaying the structural MRI images with the respective segment images in all individuals in the sample. A different approach to verify the success of the segmentation is to demonstrate the separation of the multivariate tissue intensity distribution between the voxels assigned to the vessels segment and the others. This approach, which allows visualizing the segmentation results across the whole sample at once, is presented in the Methods section.

A second feature of the segmentation is the distribution of the tissues loading on the two Gaussians of the mixture modelling bone, which we found to be reproduced with considerable consistency across the sample (labelled ‘bone 1’ and ‘bone 2’). The different intensities modelled by these two Gaussians arise from the spongious inner part of the skull, which has slightly higher intensity than the laminar portion especially in T2-weighted images. In Figure 3 these two separate loadings are shown with different colours. In the higher portions of the skull, the first sub-segment (‘bone 1’, dark blue) corresponds spatially to the laminar bone components, while the second (‘bone 2’) corresponds spatially to the diploe, which can be discerned by the higher intensity in the T2-weighted images compared to the laminar bone. There was some variability in the success in identifying this component in some subjects, however. Diploic components were identified predominantly in the frontal and superior parts of the skull, while they were less conspicuous in the lateral and posterior regions. The ‘bone 1’ component was especially represented at the base of the skull, where it was detectable in the temporal bone/mastoid process.

Figure 4 shows boxplots of the correlation between the mean grey matter signal course and that of the other tissue classes. The correlation of the bone segment was approximately the same as with the standard segmentation, averaging at approximately 0.65. The boxplots corresponding to the two bone sub-segments suggest that the high correlation can be mainly attributed to ‘bone 1’, reaching a slightly higher correlation value than the whole bone segment, while ‘bone 2’ averages at roughly 0.3. The correlations of the mean grey matter with white matter, soft tissue and ventricles replicate those of the previous analysis. These results are consistent with those of the previous analysis as far as tissue with large venous collectors (the diploe with the diploic veins, the base of the skull) is less well associated with the mean grey matter signal than the rest of the bone.

**Figure 4.**
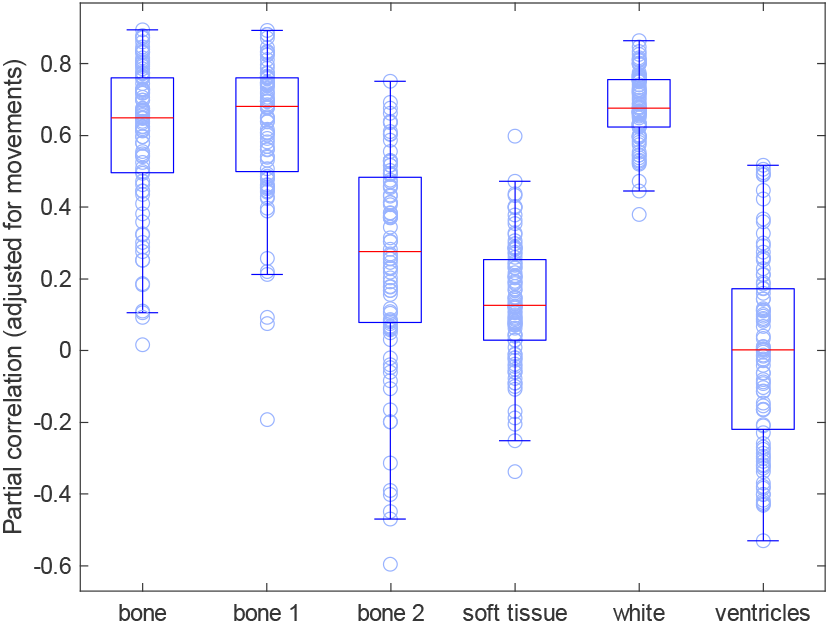
Boxplot of partial correlations between signal courses of average grey matter and bone, ‘bone 1’, ‘bone 2’, soft tissue, white matter, and ventricles, respectively. Data were adjusted for movement covariates, their first derivative, and their quadratic terms.

To explore reasons for individual differences in the degree of association between mean grey and bone signal courses, we looked at the signal course in three individuals, one with high and two with low grey matter-bone correlation values, chosen to be representative of the variety of signal characteristics we encountered.

The participant in Figure 5 is the same as already shown in Figure 1C (bottom left), where it showed much lower association between grey and bone mean signal courses than most participants when using standard segmentation (reproduced on the left). As in the analysis of Figure 1C, the association between grey and bone was low when looking at the whole bone segment. We see here that the low association had two sources: the signal from the ‘bone 2’ segment, which we have seen to come from the inner portion of this segment, and the signal from the vessels segment, which showed a marked delay relative to that of grey matter (right part of the figure, middle and bottom traces). In contrast, the signal from the ‘bone 1’ segment showed a good association with grey matter. Hence, the more specific segmentation could demonstrate a signal associated to that of grey matter in the cranial bone that was not evident using standard segmentation.

**Figure 5.**
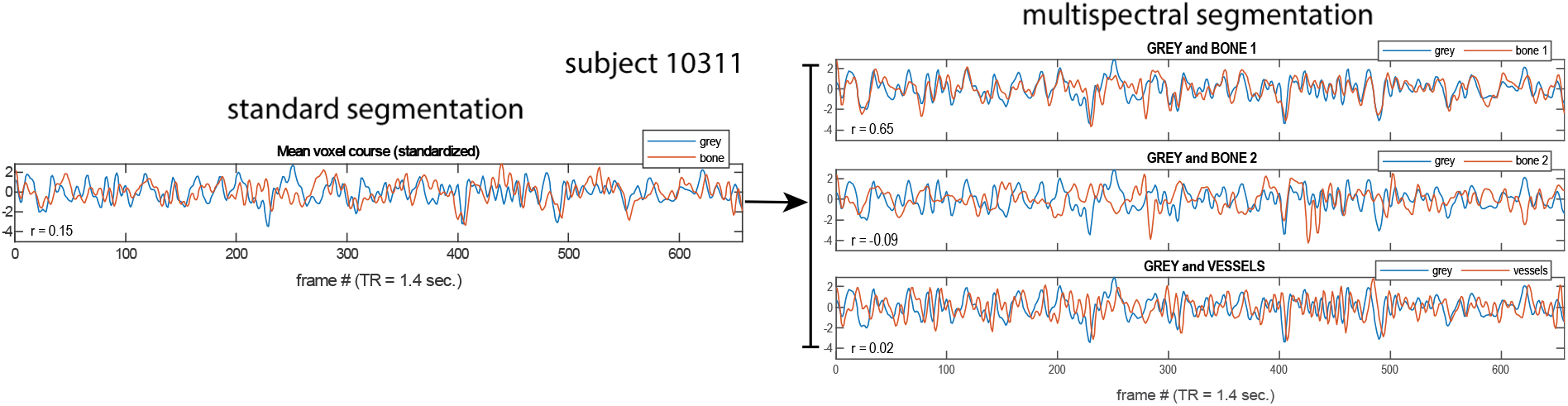
Left: mean signal course of the grey matter and cranial bone segment using standard segmentation (reproduced from Figure 1C). Right: using multispectral segmentation, the signal from bone can be further decomposed in components stemming from ‘bone 1’, ‘bone 2’, and vessels (from top to bottom). The signal was standardized in all plots.

The influence of the ‘bone 2’ segment, however, cannot explain all low correlations between bone and grey matter. As is apparent from Figure 4, there were a few participants showing low correlations between grey and bone even when using the ‘bone 1’ segment. The mean signal from the multispectral segmentation of one such participant is shown in Figure 6, left. We see here that, as in the previous participant in Figure 5, the course of the ‘bone 1’ segment appears to be similar to that of the grey matter signal. Here, however, the signal was affected by a delay like in the vessels segment, which negatively impacted the correlation coefficient.

**Figure 6.**
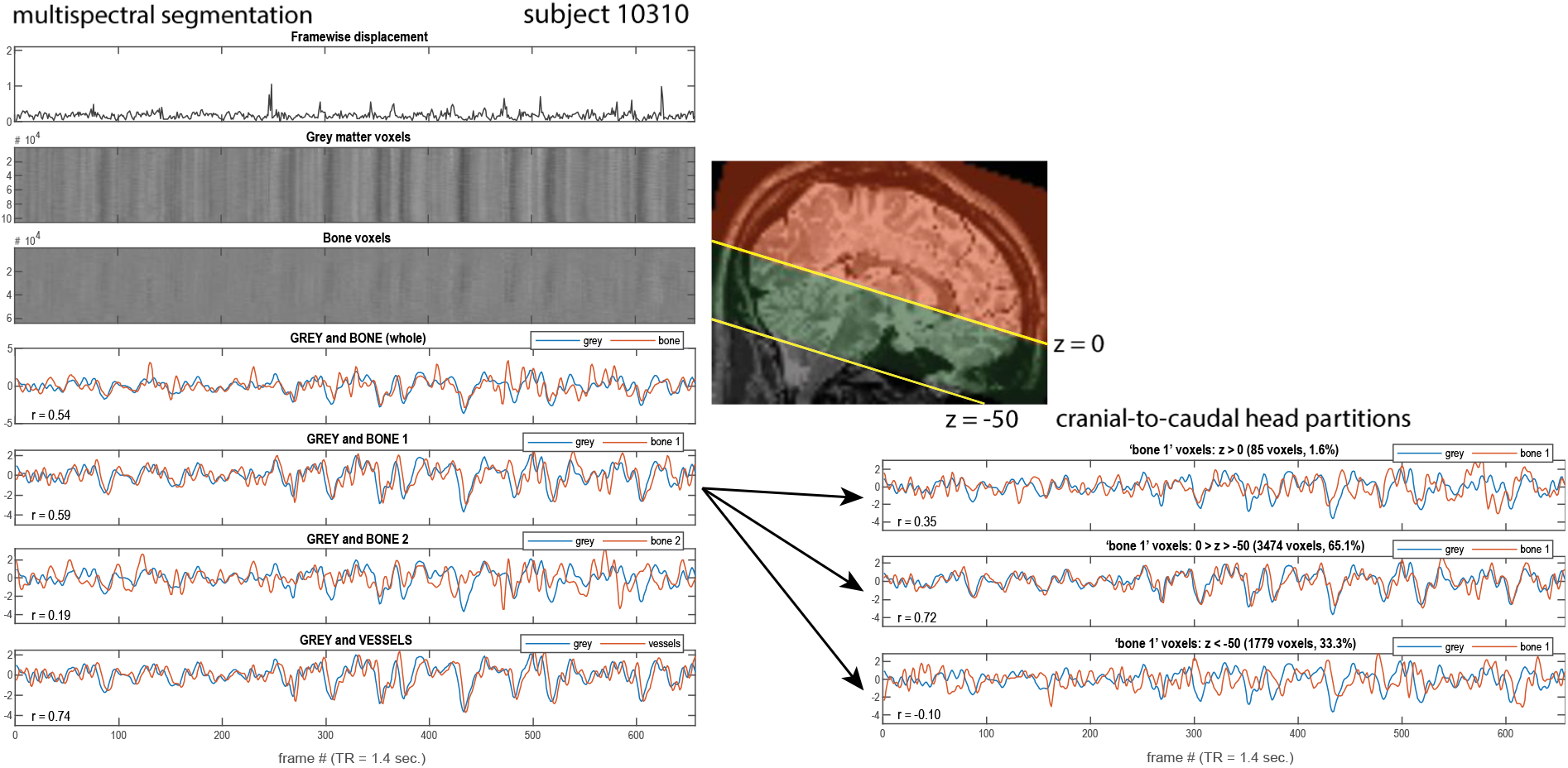
Signal course in the multispectral segmentation of subject 10310. Left, from top to bottom: the framewise displacement, the intensity of the signal in the grey matter segment, in the bone segment (both show smoothed (FWHM 8mm) data transformed to MNI space), the average signal in these two segments, the average signal of grey and the two bone sub-segments (‘bone 1’ and ‘bone 2’), the average signal of grey and vessels (no spatial smoothing, de-meaned, band-pass filtered, standardized). Right: the average signal of grey and ‘bone 1’ separated for different head partitions using masks including z > 0 (orange part in the inset), 0 > z > -50 (green part), and z < -50 (grey part), respectively. The masks were reversely transformed from MNI space into native space.

In one of the previous standard segmentation analyses, we had excluded the lower portion of the head to make sure that the association between cranial bone and grey matter did not arise from signal from large vessels at the base of the head. The result was that association was better after this exclusion (Figure 2). We therefore investigated here too the differential association of bone and grey matter in separate partitions of the head, from cranial to caudal (Figure 6, right). As shown in the brain section in the inset, segments were partitioned in the portions above *z* = 0, between *z* = 0 and *z* = −50, and below *z* = −50. The results suggest that partial failures in the segmentation may have been responsible for the low association between grey and ‘bone 1’ in this subject. In the upper portion of the head, few voxels were classified as ‘bone 1’. In the middle, where segmentation appeared to be successful, the association between bone and grey matter was good. In the bottom segment, the signal was delayed. Especially at the base of the brain and head, the differentiation between the tissue classes may be more challenging. In this individual, some venous sinuses were not segmented correctly (like S. maxillaris and S. sphenoidalis) and were partly classified as bone.

Finally, we show the results of the spectral segmentation in an individual where the association between bone and grey matter was high (Figure 7). The agreement between the signal course of bone and grey matter is good in the ‘bone 1’ segment but is also present in ‘bone 2’. The signal from the vessels segment is delayed by about two frames but follows the signal from grey matter closely. Such behaviour was observed for most subjects in the current sample. If we look at the partitioned association between these two segments, we see that, as in the previous subject, the signal from the ‘bone 1’ segment below *z* = −50 is affected by delay.

**Figure 7.**
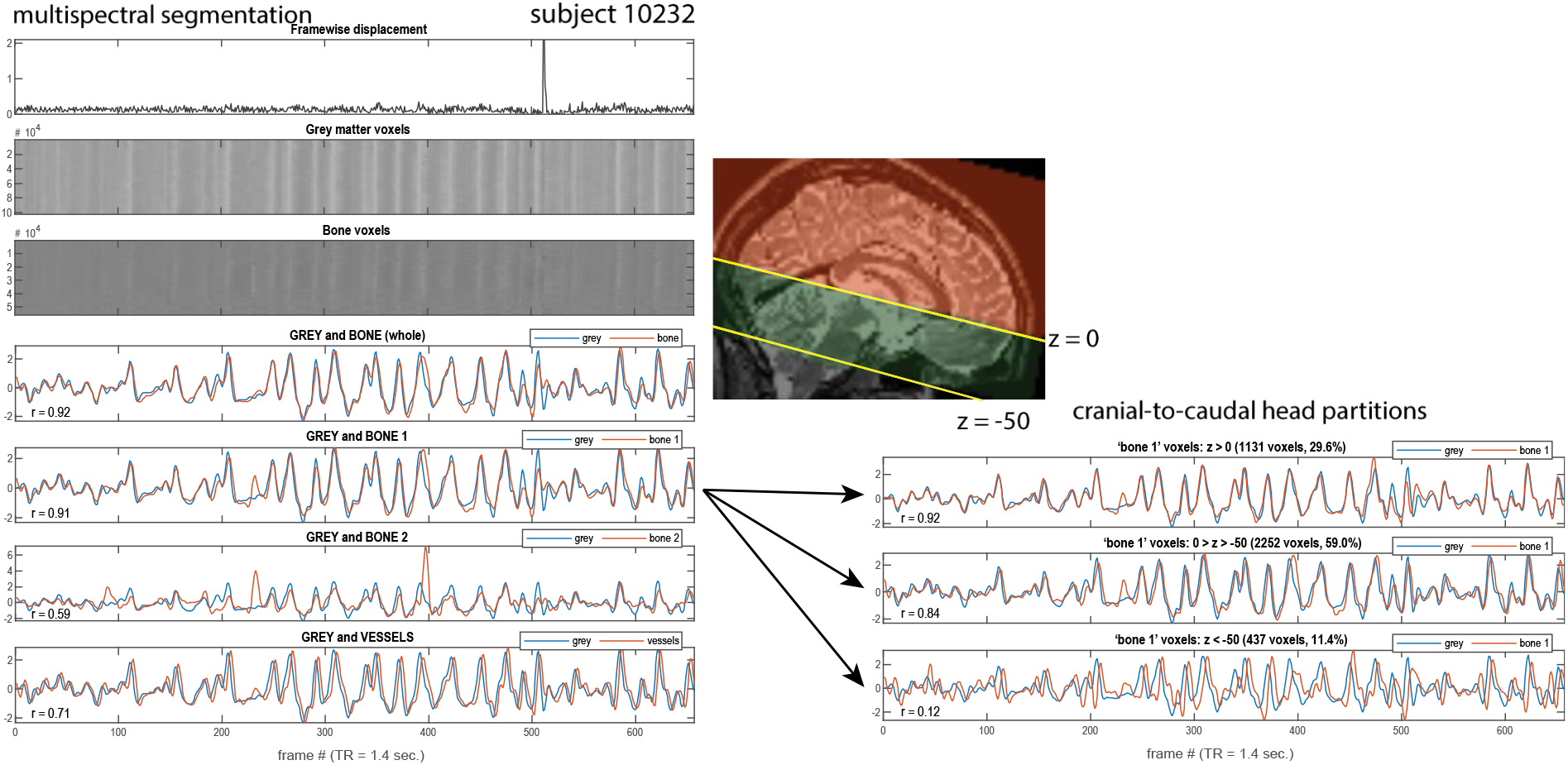
Signal course in the multispectral segmentation of subject 10232. Same conventions as in. **Figure 6**.

### Multispectral segmentation, signal delays

An aspect of previous findings was the differential delay of the mean signal of segments. Here, we looked at this issue systematically by computing lagged covariance in an AR(1) model, using the technique detailed in Mitra et al. (2014). As shown in Figure 8 (left), relative to grey matter the lag of the signal in the vessels segment amounts to about 4 sec, in ‘bone 1’ to about 1 sec. The signal in ‘bone 2’ has no average lag relative to grey matter, but this result must be viewed with caution given the large spread of the signal lag in this segment, which is much more difficult to characterize than in ‘bone 1’. Note that the lags between vessels and ‘bone 1’ and grey and ‘bone 1’ (right) are ordered across individuals (parallel lines connecting individual observations).

**Figure 8.**
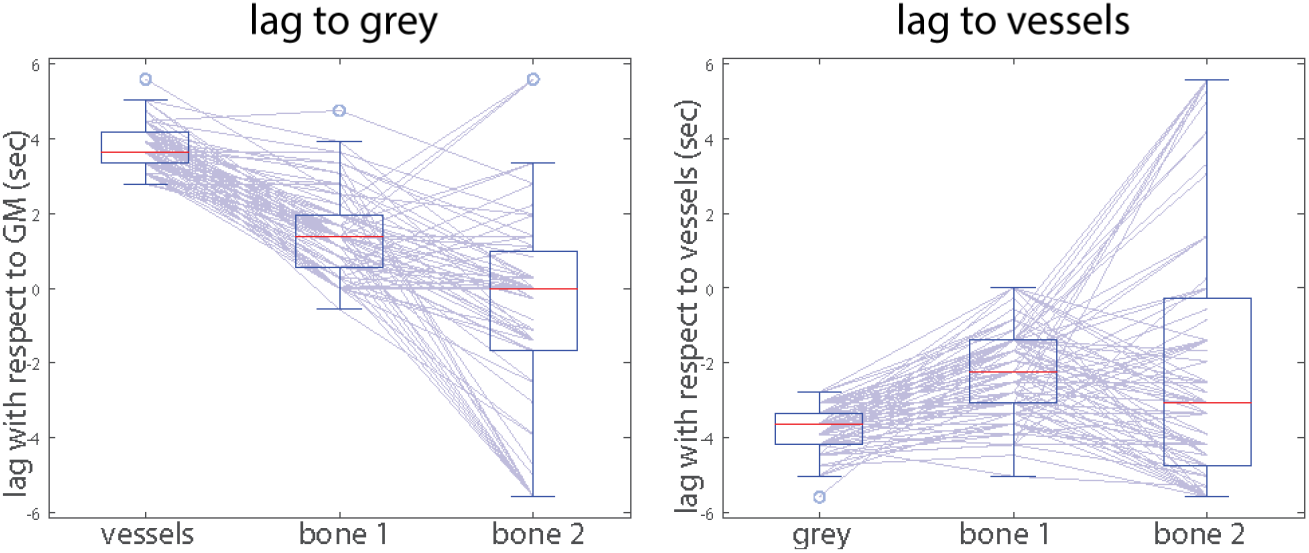
**Estimates of lags between the mean signal course relative to grey matter (left) and the vessels segment (right). The lines connect observations from the same individuals.**

### Multispectral segmentation, head motion

Around 30% of the participants exhibited considerable head motion during image acquisition. Many of these motion occurrences were accompanied by short increases in the mean grey signal, followed by a slower signal decrement, as shown in previous studies (Power 2017). One of these participants is in Figure 6, where one can see that the marked decreases of signal in grey matter were present also in the ‘bone 1’ and in the vessels segment. However, similar marked decreases may be observed also in the absence of head motion (Figure 7), as in Power (2017).

We looked at head motion again after taking the square of the framewise displacement as well as of the mean courses of the signal in the segmented data. Squaring the signal emphasizes the largest movements to the detriment of small noise-like motions, which are present throughout the course of most subjects. We found this technique useful to highlight effects of larger movements irrespective of the sign of the signal change. Note that the signal course plots differ from those of previous plots: we always show the squared framewise displacement (instead of the signal from grey) paired with the squared signal course from one segment.

Figure 9A shows a subject with head motion, resulting only in very small changes in the signal in grey matter. In bone, these signal changes are visible in ‘bone 2’, while in ‘bone 1’ only the largest movement left a mark. Hence, the signal from ‘bone 1’ followed that of grey matter while ‘bone 2’ was a conveyor of head movements. This was a common pattern in subjects with head motion. In panel B we show a subject where this pattern led to a complete dissociation between the momentary increases in the signal, present only in ‘bone 2’, while the decrements following large movements were present in both grey and in ‘bone 1’.

**Figure 9:**
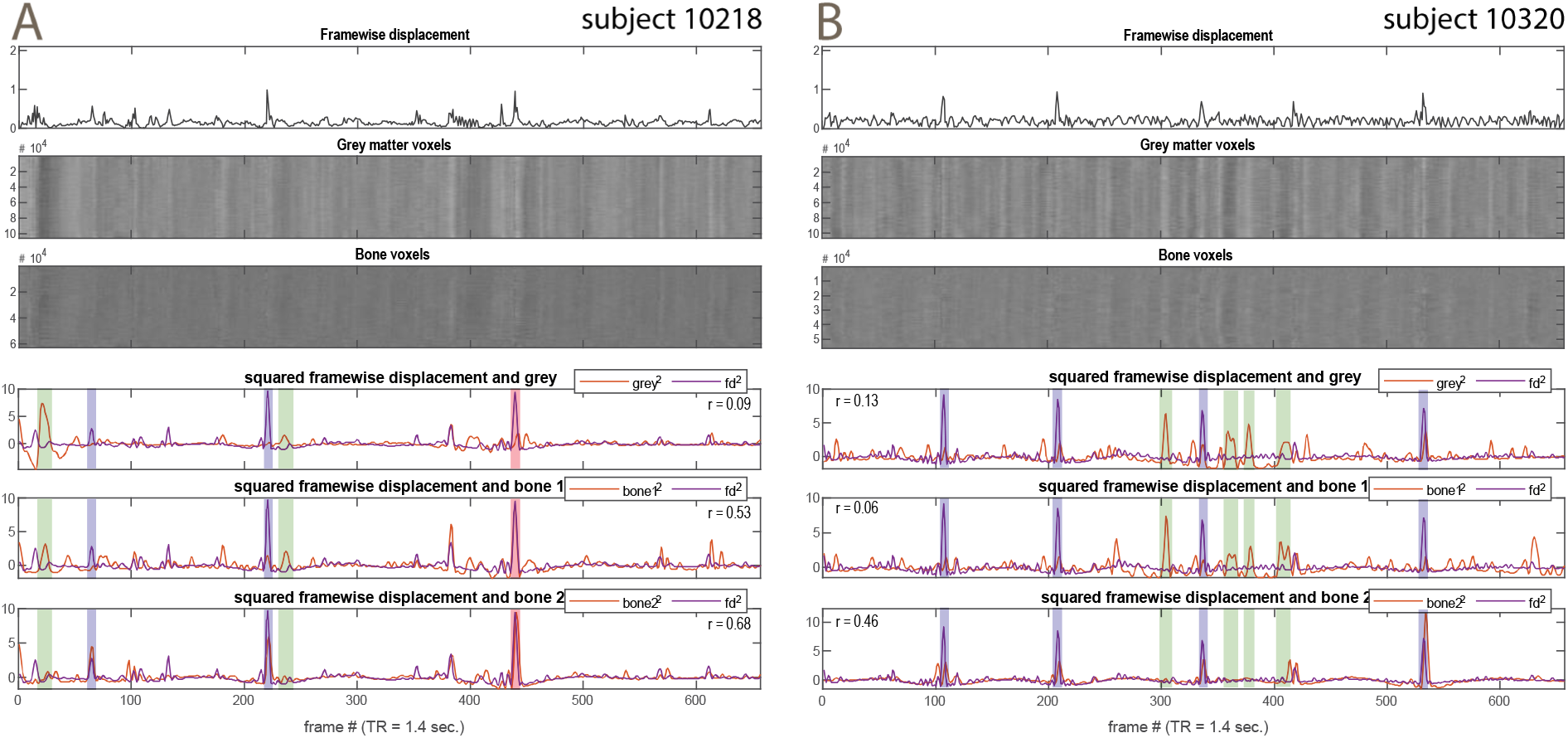
Examples for signal courses for subjects with distinct motions. The framewise displacement is shown at the top, while the three bottom plots show the squared displacement paired to squared signal courses. The blue bands show the points where large signal changes coincident with larger movements were present in the ‘bone 2’ segment, but not in grey or ‘bone 1’. The pink band shows the points where the movement signal was present in both bone segments. The green bands show the points where larger signal changes in grey were present in ‘bone 1’, but not in ‘bone 2’, which conveyed movements instead. All signals without spatial smoothing, band-pass filtered, standardized.

To characterize further the impact of motion on estimates of correlations between mean segment signal courses, we repeated the analyses of Figure 4 to compare the median correlations between bone and grey with those obtained without adjusting for movement. The median correlation between grey matter and bone in Figure 4 was 0.650, without movement correction 0.675 (paired *t* test, *t*_87_ = −1.32, *p* = 0.19). When adding frame censoring to movement correction, the median correlation was almost the same as when applying the movement correction alone (0.655). The latter value resulted from censoring all frames exhibiting a framewise displacement larger than 0.5 mm excluding also the previous frame and two frames forward (Power et al. 2012).

## Discussion

In our analysis, most subjects presented a signal from the cranial bone that was highly correlated with the mean grey matter signal. There were a few individuals where this correlation was absent, which might have been due to multiple causes, including a lower amplitude of the mean signal itself, as well as a failure to segment the cranial bone selectively. Furthermore, lower correlations may ensue in individuals where cranial bone carried signal changes attributable to head motion, as one would expect (this was especially visible when separating bone into two sub-segments). Irrespective of what the causes of these low correlations may have been, the significance of our data lies in the fact that many subjects showed high correlations between cranial bone and grey matter. The box plots of these correlations, irrespective of the method used to obtain them, show a median correlation of about 0.8, implying that half of participants had larger correlations than this. It was common to observe correlations corresponding to 80% of shared variance or more, although there would be no reason for this finding if the average grey matter signal arose mostly from neural activity.

Few studies have investigated the vascularization of cranial bone. These few data show blood supply from both extracranial and intracranial source vessels, with extracranial vessels being more involved in supplying the periosteal portion of the skull (Lammers et al. 2023). Intracranial sources are the meningeal arteries, which despite their name mainly supply the skull (Brookes and Revell 1998). In the mouse, blood supply of the skull up to the diploe by meningeal arteries has been recently demonstrated *in vivo* (Cai et al. 2019). It therefore seems plausible that the signal detected in the cranial bone was predominantly of intracranial origin. This would also explain the dissociation with soft tissue, whose association with grey matter was small, observed in our study. Before discussing what signal sources the brain and the meningeal space may have in common, we will summarize possible artefactual explanations of this commonality.

### Confounders and artefacts

In the subdural space, an important signal source was large venous sinuses, where the signal followed that of cortex but with a phase delay (Liu et al. 2017). This signal also appeared to be present in “cranial bone” when these collectors were not segregated by the segmentation. This finding may be explained by venous blood flowing from the cortex. In support of this explanation, recent high-resolution data on the spatial distribution of the BOLD signal show it to be largely localized in the veins running on the cortical surface (Kay et al. 2019). Furthermore, the vascular tree of the brain appears to affect the fMRI signal from grey matter (Tong et al. 2018; Zhong et al. 2023). However, this explanation was not applicable to the signal from bone, which showed no delay after segmenting out the contribution of vessels. If signal from the venous sinuses contaminated blood from cranial bone, we would expect this latter to be even more delayed, not less, than the signal from venous sinuses. Therefore, a different explanation seems to be required for the signal association between cranial bone and grey cortex.

Our analysis was designed to exclude other obvious explanations of the association between cranial bone and cerebral cortex, such as partial volume effects or contamination through smoothing. However, there are artefactual sources of common signal that cannot be completely excluded by our analyses. One is Gibbs artefacts arising from the reconstruction of images, giving lines paralleling high-contrast edges between tissue classes of different signal intensity. Because the signal intensity of bone is very much smaller than that of grey matter, these artefactual signals around the edges of the brain may have led to replication of the brain signal into the bone compartment. Furthermore, they may dominate the signal from bone because of its low intensity. It should be noted, however, that Gibbs artefacts are not prominent in images of low resolution such as those used in functional imaging and were not detectable here by visual inspection. Furthermore, the association between grey matter and cranial bone remained present also when one was extracted from the upper and the other from the lower part of the head, showing that spatial contiguity was not necessary to elicit the association.

Another important source of artefacts is head movements, which we know from the study of rest connectivity to be an important source of global signals (Power et al. 2012). Even small movements have been shown to affect the global signal, especially through the high frequency part of the spectrum (Satterthwaite et al. 2013). However, aggressively partialling out head movements as suggested by Satterthwaite et al. (2013) did not remove the association between cranial bone and grey matter. Furthermore, the finding that the movement-related signal was dissociable from the signal from grey matter when modelling bone with two tissue classes further argues against the global signal being prevalently due to head motion. Finally, the association between the mean cranial bone and grey matter signals was present at high levels in individuals that showed little head motion.

Another possible source of common signals is cardiac pulsations (Hu et al. 1995; Dagli et al. 1999). However, these effects are localized around large vessels (Dagli et al. 1999). Here, the association between cranial bone and cerebral cortex remained even after excluding the lower portion of the head, which contains the major portion of the arterial tree (Viviani 2016). A related source of signal changes is heart rate variability (Schmueli et al. 2007; Chang et al. 2009). However, this source has been estimated to account for about 30% of global signal variability (Power et al. 2017), which is well below the peaks of shared variance between cranial bone and grey matter found here. Similarly, although respiration has been shown to affect the fMRI signal (Wise et al. 2004; Birn et al. 2006), its contribution to the global signal may be quantified at 20-35% (Power et al. 2017; see also Hocke et al. 2016).

### Cerebrovascular reactivity, vascular self-regulation, and other forms of vascular activity

While none of these well-known cardiovascular sources of “physiological noise” may alone explain the high association between the signal from cranial bone and grey matter reported here, there is increasing evidence that they may be part of the physiology of the cardiovascular system and of the interrelated mechanisms that together regulate the arterial supply of the brain. Hence, when acting in concert, their effects may be compounded. The vascular (as opposed to neurogenic) origin of the global fMRI signal is a controversial and hotly debated issue (Liu 2013; Abdelkarim et al. 2019; Tong et al. 2019a; Bright et al. 2020; Colenbier et al. 2020; Chen et al. 2020; Chen and Gauthier 2021). Part of this debate concerns cerebrovascular reactivity, i.e. the vascular response to vasodilator stimuli such as increased CO_2_ concentration, but the same mechanisms may be involved in systemic physiology. For example, respiratory activity is associated with head motion and changes in CO_2_ partial volume, which has vasodilatory effects (Wise et al. 2004). Power et al. (2017) have reported that conspicuous head movements were associated with respiratory events and variations in heart rate and were typically followed by global signal decrements lasting 6-12 sec. that may be explained by the variations in partial CO_2_ pressure associated with these events.

The events described by Power et al. (2017), and the associated decrease in global signal, were clearly visible in the signal from cranial bone in our analyses, suggesting that some of the involved mechanisms of vascular regulation, such as the increased CO_2_ partial pressure, may be shared between the brain and cranial bone. However, CO_2_ reactivity due to respiratory events seems not to explain all fluctuations in the global signal. As noted by Power et al. (2017), not all fluctuations of the global signal were accompanied by respiratory events, apparent through conspicuous head movements.

Considerable data have been presented in the literature suggesting that vascular activity not arising from the neurovascular coupling provides an important portion of the global signal. The vascular origin of the signal is shown by the existence of spatial-temporal patterns that travel with the blood, from the arteries through the brain to large venous sinuses (Tong and Frederick 2010; Tong et al. 2019b; for a review, see Tong et al. 2019a). Recently, using an information-theoretic approach, Colenbier et al. (2020) showed that vascular signal (arising from a mixture of arterial and venous tissue) not only was strongly associated with the global signal in some individuals, but also contributed to intrinsic connectivity. The extent of the association showed a skewed distribution with a median around 0.8-0.9, as in our data. These findings are consistent with the prominent role of a vascular signal arising simultaneously in the cranial bone and grey matter, as reported here.

### Neurogenic regulation of vascular tone

There is evidence that vascular tone and flow may be under neurogenic control. Quantitative studies of cerebral flow with PET or arterial spin labelling show large-scale changes in perfusion that may be mediated by receptors acting on vessel tone (for reviews, see Hamel 2006; Viviani et al. 2014). The evidence is particularly good for cholinergic control from the nucleus of Meynert through its cholinergic projections (Sato et al. 2001; Liu et al. 2018; Turchi et al. 2018). Cholinergic agents produce marked vasodilation also in meningeal vessels (Levy et al. 2019), but there is no evidence that central cholinergic projections from the nucleus of Meynert, which affect blood flow in cerebral arteries, exit the brain parenchyma (Adachi et al. 1992), making this an unlikely mechanism to explain our findings.

Sensitivity to cholinergic agents in meningeal vessels is commonly interpreted as evidence for autonomic control, even if the role of parasympathetic activity in vasodilation of meningeal vessels is still debated (Coles et al. 2017; see also Kovacs et al. 2004). Classic studies have also provided evidence for sympathetic control of the cerebral venous system (Schmidek et al. 1985). While principally focused on cerebral vessels, these studies have investigated the effect of the stimulation of the sympathetic cervical nerve, which also innervates meningeal tissue. In summary, considerable empirical evidence exists for an effect of autonomic activity of both cerebral and meningeal vessels that may explain the commonality of the course of the mean signal in the tissue classes observed here, in addition to the mechanisms of cerebrovascular reactivity mentioned above.

The possible vascular origin of a considerable portion of the global signal does not warrant the conclusion that it constitutes “noise”. Apart from the observations from connectivity studies finding individual differences that disappear after adjusting for this signal (Gotts et al. 2013; Hahamy et al. 2014), the possibility of a role of autonomic regulation implies that this signal, or parts of it, may be themselves under central control. For example, the existence of autonomic modulation during the appraisal of stimuli, especially those conveying emotional valence and arousal, is a well-established finding in emotion research (Lang and Bradley 2010). Several researchers are actively researching the role of arousal in determining global properties of the fMRI signal (Wong et al. 2013; Raut et al. 2021). Martin et al. (2021) have reviewed the evidence that states other than arousal, such as emotional activation, may affect estimates of resting state connectivity. These findings may explain the existence of different correlates of the global signal, as a variable such as arousal may affect the global signal through different mechanisms simultaneously.

### Concluding remarks

The large association between the mean signal from cranial bone and grey matter provides an avenue for future research aiming at clarifying the extent to which the global signal is of vascular nature. Addressing this issue is important for several reasons. One reason is clarifying the extent of deactivations in task-based fMRI (Viviani et al. 2020; Labek et al. 2023). As has been noted, adjustment for the grey matter signal equalizes task activations and deactivations (Zarahn et al. 1997). For this reason, task deactivations may be thought to be artefactually too high after adjustment for physiological noise, whether obtained by adjusting for the grey matter signal or other intracranial sources, as all these signals are reciprocally correlated. A large vascular component in the global signal would imply that the large task deactivations obtained after adjusting for bone and other intracranial signal sources (other than grey matter) are not an artefact of adjusting for “physiological noise”.

Another issue of great importance is disentangling different sources of individual differences in neuroimaging data. Models of the global signal that attribute it to autonomic or arousal processes imply that individual differences in this signal may have their origin in central regulation but may be misinterpreted as arising directly from the neural signal of cortex. This justifies paying renewed attention to the possible constituents of the global signal in future task-based functional imaging studies of individual differences.

## Methods

### Sample and data acquisition

The sample consisted of 88 participants (15 females, age range 20-50) taken from the MPI-Leipzig Mind-Brain-Body dataset (Mendes et al. 2019). The subjects were chosen from a bigger pool according to the availability of structural images, and after excluding individuals of age 50 or older to avoid the possible confounder of systematic differences in the effectiveness of the segmentation. In order to perform the intended analyses, multiple modalities of high-resolution structural images were required, i.e., T2-weighted image (TR=3200 ms, TE=409 ms, flip angle 120°), high-resolution 3D FLAIR (TR=5000 ms, TE=395 ms, flip angle 120°), two MP2RAGE images (TR=5000 ms, TE=2.92 ms, TI=700 ms and 2500 ms, flip angle 4° and 5°, respectively). The functional images were acquired during a 15-minute eyes-open resting-state session (TR=1.4 s, TE=30 ms). The study was conducted with a 3T Siemens Verio scanner. Additional details on image acquisition may be found in the dataset description publication (Mendes et al. 2019).

### Standard segmentation

The presented results were obtained with two different pre-processing pipelines – both based on the functions of the SPM12 package (Ashburner and Friston 2005). The first one uses the default settings given in the software (‘Standard segmentation’ in the Results). The functional images were realigned by rigid-body transformations yielding their mean image together with the movement parameters. Mixtures of Gaussians were applied to model the intensity distributions based on spatial priors for the respective tissue at the voxel coordinates. The mean functional image was taken as the only input channel for segmentation of the six tissue types, which are defined in the tissue probability maps of the software package: grey matter (GM), white matter (WM), cerebrospinal fluid (CSF), bone, soft tissue, and background.

To make sure that the data used in the Results did not contain signal from adjacent segments, we applied an erosion of 1 voxel for grey, 2 for bone, and 2 for soft tissue to the respective segments. We limited the erosion of the grey segment to 1 voxel because the large surface of this segment causes large tissue losses with erosions of 2 voxels or more. In the data registered to MNI space (Figure 1B-C), which were resampled to a standard 2mm isotropic voxel space, this produces a separation of about 6mm between bone and grey matter. The signal from CSF was masked by a ventricle mask from the Harvard-Oxford Cortical and Subcortical Atlas (https://fsl.fmrib.ox.ac.uk/fsl/fslwiki/Atlases). The signal was band-pass filtered between 0.1 and 0.0078 Hz. In the plots of the results, partial correlations were computed to partial out the contribution of the movement parameters from realignment, their first derivative, and their quadratic terms (24 parameters in all).

The same procedures were applied to native data for the results of Figure 2, where voxel size was 2.3mm, giving slightly larger margins between the segments used to estimate mean signal courses. The masks used for the results of Figure 2 were first created in MNI space. The affine transformation computed by the SPM12 package to register volume was reversed and applied to the masks, giving masks in native space.

### Multispectral segmentation

The second pre-processing pipeline used a multivariate approach to improve the accuracy of tissue assignments (‘Multispectral segmentation’ in the Results section). In multispectral segmentation, images of different modalities are used to assign posterior probabilities of individual voxels to belong to a tissue class (Lambert et al. 2013, Viviani et al. 2017). The posterior probabilities are computed after fitting a mixture of Gaussians to the density of image intensities, which has the dimensionality of the number of input modalities used in the segmentation. As priors, tissue density given by the expected location of the tissue classes in MNI space are used (in the SPM12 approach, segmentation internally computes a transformation to MNI space to register the prior to the voxel intensity observed in the native images, outputting posterior probabilities in both native and registered spaces). Because MRI modalities provide different tissue contrasts, using more than one modality allows one to better identify a tissue class using the information from the modality where the contrast is high. This is because the modality where the contrast is high causes the multivariate density to differentiate between the tissue classes. Here, we used this approach to increase the number of tissue classes that were segmented, adding one tissue class for brain vessels and subdural sinuses, and identifying two sub-segments in the bone tissue class.

Besides the mean functional image, four structural images were set as input channels for the segmentation: high-resolution FLAIR, MP2RAGE (two images with different inversion times), and T2w. All structural images were co-registered to the mean functional image before starting the segmentation procedure. Additionally, an extended set of priors (TPMs) was applied, which includes a separate tissue class for vessels (VES) based on expected density of vessel signal (Viviani 2016). To account for different signal intensities belonging to the same tissue class, more than one Gaussian can be used to model a certain tissue type (Viviani et al. 2017). The number of Gaussians for each tissue class, leading to the most accurate results, was defined after testing several combinations: GM 2, WM 2, CSF 2, bone 2, soft tissue 4, VES 2, background 4. Tissue allocation was evaluated by visual inspection of the respective segment images when overlaid on the source images (i.e., input channels). It was found that better results were obtained when at least two Gaussians were used for each tissue class. Otherwise, single-Gaussian tissues were slightly compromised by their surroundings. We verified that two Gaussians were adequately modelling the bone tissue class, giving repeatable mixture models across the sample. Figure 10 shows the outcome of the mixture model. The real densities are in five-dimensional space. In the Figure, we plotted the densities using two dimensions in each plot; each plot represents the same data using different dimensions.

**Figure 10.**
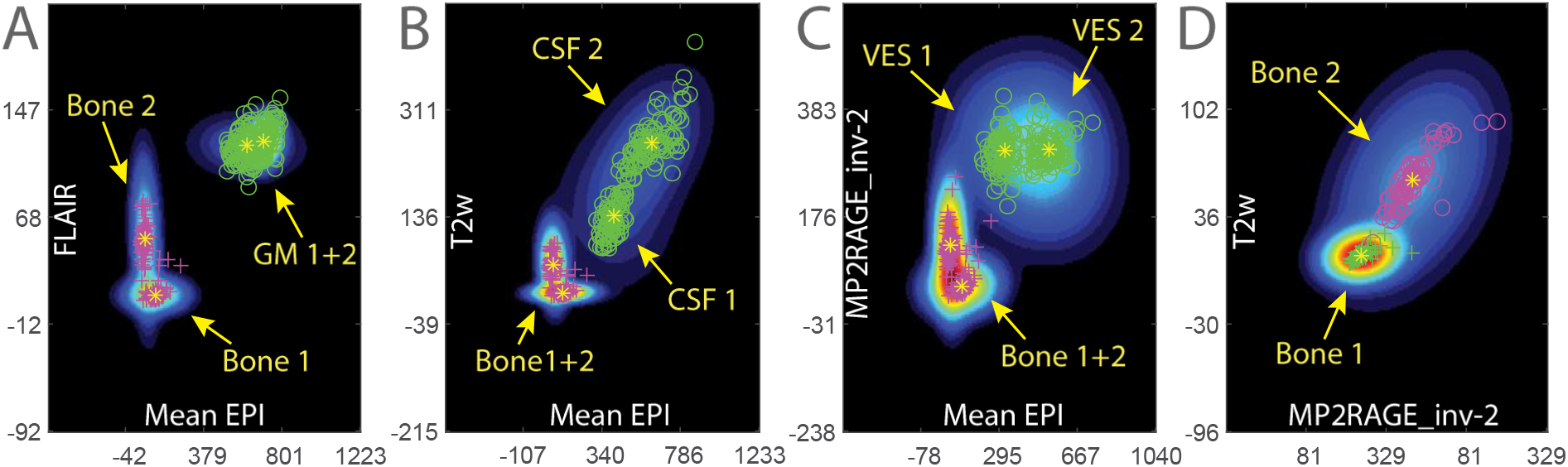
Tissue class densities estimated from the whole sample (colour gradient from shades of blue to red), showing the centres of mass of individual participants. A: densities of bone and grey matter (GM), B: bone and cerebrospinal fluid (CSF), C: bone and vessels (VES), and D: the separation between the two bone sub-segments.

Our concern in this segmentation was to separate bone from grey matter and the vessels segment (to avoid as much as possible confounders in assigning the mean signal to the wrong tissue class), and to explore the extent to which it was possible to further divide the bone segment into two tissue sub-classes. Figure 10 shows the modelled tissue class density (colour gradient from dark blue to red), and individual points represent the density peaks in the volumes from individual participants. Figure 10A-C shows these estimated quantities for the bone Gaussians with respect to GM, CSF, and VES, respectively. The success of the segmentation can be assessed in two respects. The first is the consistency with which the peak intensities were assigned to the same density across the whole sample. This criterion was satisfied in all participants, as the plots show. The second is the extent to which the densities themselves were separate, as this represents the capacity of the contrast to differentiate between tissues. All tissue classes show a clear separation when plotted in the modality with the appropriate contrast. In the case of bone and CSF, both the EPI modality and T2-weighted images contributed to the separation, because the density in multivariate space produces a region of space of low posterior probability for both that was not present in the projection in the individual dimensions (Figure 10B). Because joint probabilities are multiplicative, the existence of low probabilities in one direction of space is enough to take the posterior down to low values, irrespective of the probabilities in other directions.

Figure 10D shows the separation of the two bone sub-segments in structural images (in Figure 10A-C one can see that the bone is uniformly of similarly low intensity in the EPI modality). In both T1- and T2-weighted images, there is a bone Gaussian peaking at higher intensities (‘bone 2’), corresponding to the internal part of bone (including internal porous bone and the diploe) as shown in Figure 3. However, the separation between bone sub-segments is more difficult than that between bone and other tissue classes. One can see that there is some overlap between the Gaussians, which can also be attributed to insufficient image resolution with respect to the dimensions of the related tissues. One can also see that one of the Gaussians has very localized peaks and density (‘bone 1’), indicating that signal with the required intensity was identifiable with more consistency across the sample than in the other Gaussian (‘bone 2’).

### Statistical analyses

Statistical computations were conducted in MATLAB (R2023a). Partial correlations were determined with the function *partialcorr*. The calculation of lagged correlations followed the method described by Mitra et al. (2014). First, all signal courses were shifted to zero-mean and scaled to unit variance. Next, the temporal resolution of the time-series was increased by up-sampling (factor 5) using the spline interpolation method of the function *interp1*. The movement-adjusted partial correlation coefficient between the up-sampled data of two tissue classes was maximized with respect to the time lag. The range of time deviation was restricted to four image frames, equalling 5.6 s, in both directions. In this way, time lags and associated correlation coefficients were obtained in reference to grey matter and vessels, respectively. By definition, negative lag values denote a preceding signal relative to the reference, while positive values indicate a retardation.

## Data and software availability

Data of this study are publicly available at https://openneuro.org/datasets/ds000221. The code of the analyses is available at https://git.uibk.ac.at/c7201031/ztr2303.git.

## Acknowledgments

The dataset used in this study was obtained from the OpenfMRI database. Its accession number is ds000221.

A preliminary version of this work was presented at the CogBases workshop (Paris, Institut Pasteur, 10-11 October 2023). This work was supported by an ERA-PERMED grant (project ArtiPro) of the FWF Austrian Science Fund (grant number I 5903) to Roberto Viviani. The authors declare no conflict of interest.

## Notes

https://git.uibk.ac.at/c7201031/ztr2303.git

### Competing Interest Statement

The authors have declared no competing interest.

### Summary of Updates

Analysis of time lags of tissue classes added. Minor revisions text and figures.

